# NanoHIV-DR: a Nextflow workflow for consensus and read level HIV-1 drug resistance profiling from Oxford Nanopore sequencing

**DOI:** 10.64898/2026.11.20.768148

**Authors:** Daniel Bugembe Lule, Deogratius Ssemwanga, Pontiano Kaleebu, Damien C. Tully

## Abstract

Oxford Nanopore sequencing offers a rapid and portable approach for HIV-1 drug resistance (HIVDR) genotyping, but analysis of the resulting long-read data requires multiple bioinformatics steps and currently available nanopore workflows are limited. We developed NanoHIV-DR, a Nextflow-based workflow that streamlines the analysis of Oxford Nanopore sequencing data and generation of HIVDR reports. NanoHIV-DR processes raw POD5 files through basecalling, demultiplexing, host-read removal, quality filtering, read alignment and consensus generation, before identifying resistance-associated mutations and interpreting predicted drug susceptibility. The workflow provides two complementary approaches to HIVDR analysis: consensus-based resistance interpretation using Sierra and read-level detection of resistance-associated variants, enabling mutations below the consensus level to be identified and quantified. Outputs are integrated with clinical metadata to generate human-readable HIVDR reports. NanoHIV-DR provides a reproducible and accessible workflow for Nanopore-based HIVDR analysis with reduced bioinformatics expertise and manual processing for translating long-read sequencing data into interpretable resistance profiles.

**Availability and implementation:** NanoHIV-DR is implemented as a Nextflow workflow and is freely available at https://github.com/Tully-Lab/NanoHIV-DR. The current release (v1.0.0) is archived and available at https://doi.org/10.5281/zenodo.23402055.

## 1. Introduction

HIV-1 drug resistance (HIVDR) remains a major challenge to maintaining effective antiretroviral therapy, creating a continuing need for reliable and accessible resistance genotyping (Landovitz, et al., 2023). Conventional HIVDR genotyping has largely relied on Sanger sequencing, which typically detects variants present at approximately 20–30% or higher of the viral population (Avila-Rios, et al., 2020; Bonsall, et al., 2020; Kantor and Gupta, 2023). Next-generation sequencing (NGS) offers greater sensitivity for minority variants, while long-read Oxford Nanopore Technologies (ONT) sequencing can generate high-depth reads spanning large genomic regions and potentially resolve linked resistance-associated mutations and viral haplotypes (Lule, et al., 2025).

Translating raw ONT data into interpretable HIVDR profiles, however, requires multiple computational steps, including basecalling, demultiplexing, quality control, host-read removal, sequence alignment, variant detection and resistance interpretation. Although tools for Nanopore-based HIV analysis have emerged, including Nano-RECall, NanoHIV and ClusterV-Web (Delaney, et al., 2023; Su, et al., 2024; Wright, et al., 2021), their applications and analytical approaches differ, and accessible, reproducible workflows that integrate raw ONT processing with both consensus- and read-level HIVDR analysis remain limited.

Here, we present NanoHIV-DR, an open-source Nextflow workflow for HIVDR profiling from Oxford Nanopore sequencing data. NanoHIV-DR integrates sequence processing with consensus-based Stanford HIVdb interpretation and read-level detection and quantification of resistance-associated variants. These results are combined with sample metadata to generate human-readable HIVDR reports, providing a reproducible workflow from raw Nanopore sequencing data to interpretable resistance profiles.

## 2. Methods

### 2.1 Workflow implementation and sequence processing

NanoHIV-DR was implemented in Nextflow as a modular workflow for processing Oxford Nanopore Technologies (ONT) HIV-1 sequencing data and generating drug-resistance profiles (**Fig. 1**). The workflow accepts raw POD5 files or previously basecalled FASTQ files and currently targets the protease (PR) and reverse transcriptase (RT) regions of pol. Software dependencies are containerised using Docker, while analysis parameters and computational resources can be configured through Nextflow (Di Tommaso, et al., 2017). For raw POD5 input, basecalling and demultiplexing are performed using Dorado, with the sequencing model and barcoding kit specified by the user. Potential human-derived reads are optionally removed using SanitizeMe against the human_g1k_v37 reference, followed by read-quality and length filtering using NanoQ (Steinig and Coin, 2022). Quality-filtered reads are then analysed in parallel using consensus- and read-level workflows. For consensus analysis, reads are aligned against the HIV-1 HXB2 pol reference using Minimap2 (Li, 2018), and a depth-aware majority consensus is generated using iVar (Grubaugh, et al., 2019). Default thresholds require a minimum base quality of 15, an allele frequency of 0.5, and a sequencing depth of 20 reads, although these parameters are user-configurable. Samples that fail to generate an adequate consensus are recorded before downstream analysis.

**Figure 1.**
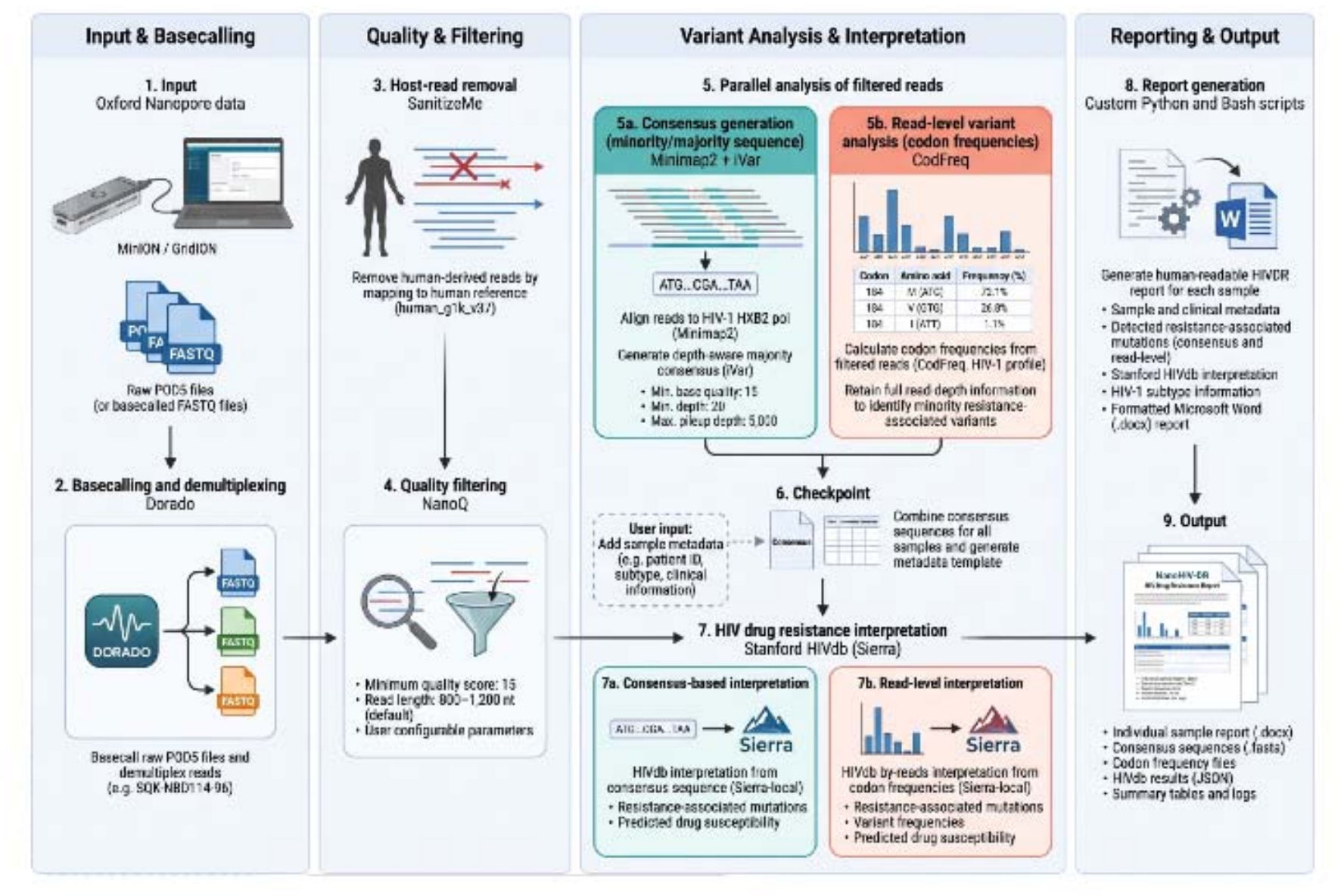
NanoHIV-DR workflow for Nanopore-based HIV-1 drug resistance genotyping and reporting. The workflow comprises four stages: input and basecalling, quality control and filtering, variant analysis and interpretation and automated reporting. Human-derived reads are removed using SanitizeMe by mapping against the hg19 human reference genome, followed by NanoQ quality filtering using user-configurable parameters. Filtered reads are analysed in parallel using two complementary approaches: (i) consensus generation with Minimap2 and iVar to produce depth-aware majority consensus sequences, and (ii) read-level variant analysis with CodFreq to quantify codon frequencies while retaining information on within-sample minority variants. At a user checkpoint, sample metadata are incorporated and consensus sequences are combined with the metadata template. HIV-1 drug-resistance interpretation is then performed using Stanford HIVdb (Sierra) for both consensus sequences and read-level codon frequencies, providing resistance-associated mutations and predicted drug susceptibility, with the read-level analysis additionally reporting variant frequencies. Finally, custom Python and Bash scripts integrate sequence, resistance, subtype, and read-level variant results to generate human-readable HIVDR reports and associated output files, including consensus FASTA sequences, codon-frequency files, HIVdb results, summary tables, and logs. The figure was created in BioRender. Tully, D. (2026) https://BioRender.com/4utcpfk.

### 2.2 HIVDR analysis

To provide complementary representations of HIVDR, NanoHIV-DR performs resistance analysis at both consensus and read levels. Consensus sequences are interpreted using the Stanford HIV Drug Resistance Database (HIVdb) algorithm through Sierra (Liu and Shafer, 2006), generating resistance-associated mutation calls and predicted antiretroviral susceptibility. In parallel, quality-filtered reads are analysed using CodFreq to determine codon frequencies while retaining within-sample variation that is lost during majority-consensus generation. CodFreq results are interpreted using the Stanford HIVdb by-reads workflow, enabling resistance-associated variants occurring below the consensus threshold to be identified and quantified. HIVdb results are retained in JSON format, together with consensus FASTA and codon-frequency files, for downstream analysis.

### 2.3 Metadata integration, reporting and reproducibility

Sequence processing and report generation are separated by a user checkpoint at which sample-associated metadata can be added. Before reporting, NanoHIV-DR verifies correspondence between identifiers in the metadata and consensus sequence files to reduce the risk of incorrect sample assignment. Custom Python and Bash scripts then integrate metadata, subtype, consensus-derived HIVdb results, and read-level resistance information to generate individual human-readable HIVDR reports. Nextflow coordinates workflow execution and provides process-level logging, caching, and resumption of interrupted analyses. Individual failed processes are retried twice before the affected sample is excluded, allowing analysis of the remaining samples to continue. Intermediate and final outputs, including consensus sequences, codon-frequency data, HIVdb results, summary files, and workflow logs, are retained to support reproducibility and downstream analysis.

## 3. Results and application

### 3.1 Workflow execution and HIVDR reporting

NanoHIV-DR provides an integrated workflow for processing Oxford Nanopore HIV-1 sequencing data from raw POD5 files or basecalled FASTQ reads through to drug-resistance interpretation and reporting (**Fig. 1**). The workflow generates majority consensus sequences for Stanford HIVdb interpretation alongside codon-frequency data for read-level resistance analysis, enabling the identification of resistance-associated variants occurring below the consensus threshold.

NanoHIV-DR was evaluated using 96 HIV-1 sequencing samples (Lule, et al., 2026) all of which successfully completed sequence processing and generated consensus and read-level HIVdb results. For each sample, the workflow combines resistance predictions, mutation frequencies, sequencing depth and sample metadata into an individual report (**Fig. 2**), while retaining consensus sequences, codon-frequency data, HIVdb JSON results and intermediate files for downstream analysis. Nextflow caching and sample-level error handling support workflow recovery without repeating successfully completed processes.

**Figure 2.**
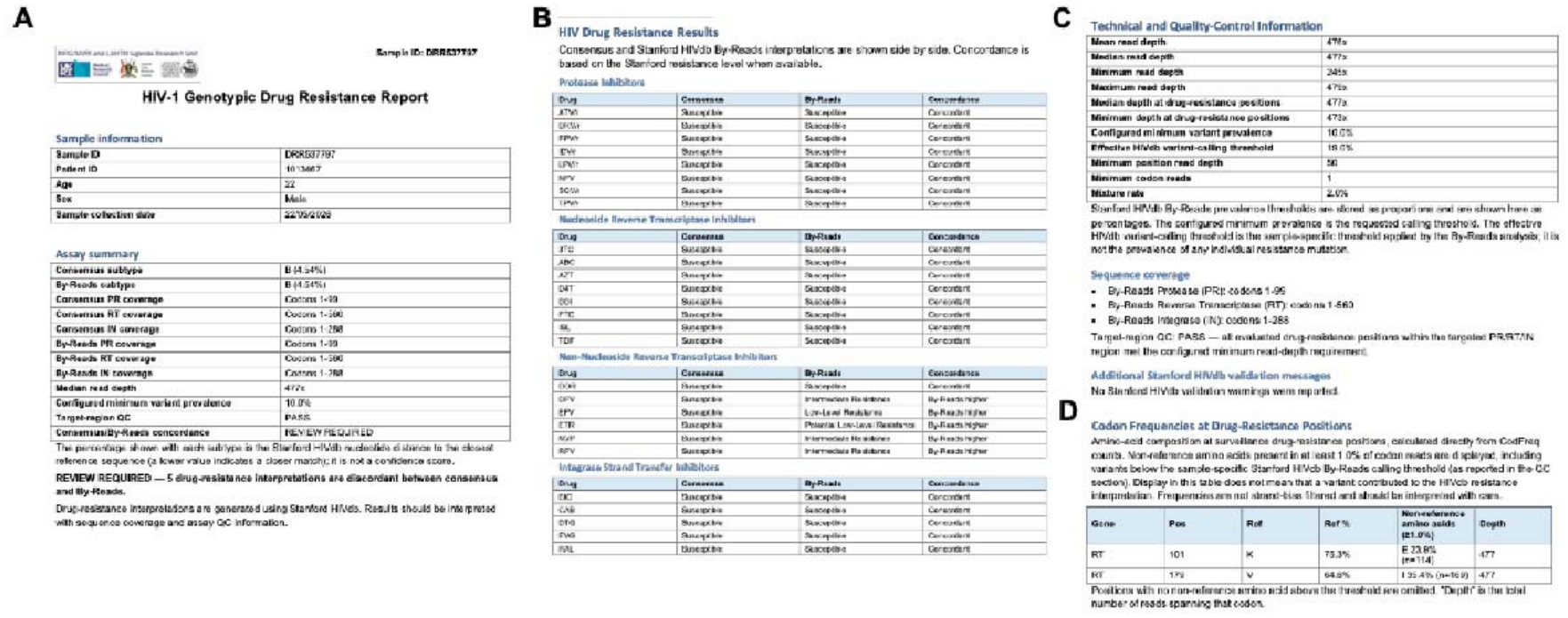
Representative clinical reporting and HIV-1 drug resistance analysis generated by NanoHIV-DR. **(A)** Example clinical report for sample DRR537797 from Ode et al. that summarises sample information, sequencing coverage and quality-control metrics. **(B)** Side-by-side comparison of Stanford HIVdb resistance interpretations across four antiretroviral drug classes. Consensus and read-based analyses were concordant for protease inhibitors, nucleoside reverse transcriptase inhibitors and integrase strand transfer inhibitors, but differed for five non-nucleoside reverse transcriptase inhibitors (NNRTIs). **(C)** Sequencing depth, variant-calling thresholds and quality-control metrics, demonstrating a median read depth of 477×. **(D)** Codon-frequency analysis identifying minority amino-acid variants at reverse transcriptase positions 101 and 179. K101E was detected in 23.9% (114/477) of codon reads, while V179I was detected in 35.4% (169/477). K101E was absent from the majority consensus but included in the read-based HIVdb interpretation as the mixed amino-acid designation K101KE.

### 3.2 Application to independently generated Nanopore data

To assess portability, NanoHIV-DR was applied to the publicly available ONT dataset of Ode et al. (Ode, et al., 2024), comprising 106 clinical HIV-1 samples and two control strains. Of these, all datasets completed quality filtering and alignment and generated consensus sequences with codon-frequency results produced for HIVdb interpretation.

This demonstrated the ability to process independently generated ONT sequencing data without modifying the underlying workflow. The analysis assessed workflow compatibility rather than independent HIVDR accuracy, as matched sample-level reference resistance profiles were unavailable.

### 3.3 Computational performance

Computational performance was assessed using a 96-sample run from basecalled FASTQ files on a MacBook Pro (Apple M1 Max, 10-core CPU, 32 GB RAM) with Docker. Sequence processing completed in approximately 2 h 52 min (22.2 CPU-hours), with codon-frequency estimation and host-read depletion accounting for approximately 85% of total CPU consumption (**Table 1**). Host-read depletion had the highest observed memory requirement (12 GB), while other processes required less than 3 GB.

**Table 1.** Per-process runtime and peak memory for Stage 1 (consensus generation), profiled on a 96-sample run from basecalled FASTQ.

| **Process** | **Mean time per sample** | **Mean CPU utilisation** | **Peak memory** |
|---|---|---|---|
| Codon-frequency estimation (CodFreq) | 53 s | 112% | 2.8 GB |
| Host-read depletion (SanitizeMe) | 46 s | 274% | 12.0 GB |
| iVar consensus | 9 s | 125% | 0.7 GB |
| Reference alignment (Minimap2) | 1 s | 406% | 0.3 GB |
| Quality filtering (NanoQ) | <1 s | 82% | <0.1 GB |

HIVdb interpretation and report generation required an additional 2.4 minutes, giving a total runtime of approximately three hours for 96 samples. These measurements exclude Dorado basecalling and were obtained under linux/amd64 emulation on Apple Silicon. HIVdb interpretation requires internet connectivity, and its runtime may vary with network conditions and server availability.

## 4. Discussion

Although several bioinformatics tools have been developed for HIV-1 analysis using Oxford Nanopore Technologies (ONT) sequencing, relatively few provide integrated workflows specifically designed for HIV-1 drug resistance (HIVDR) genotyping. To address this need, we developed NanoHIV-DR, a reproducible workflow that converts ONT sequencing data into interpretable HIVDR results. By integrating workflow orchestration, containerised software dependencies, sequence processing and resistance interpretation, NanoHIV-DR reduces the need for multiple independent computational steps and manual intervention. The workflow also combines sequence-derived results with sample metadata to generate sample-specific reports while retaining machine-readable outputs for downstream analysis.

One important feature of NanoHIV-DR is its complementary representation of HIVDR at the consensus and read levels. Consensus sequences enable established resistance interpretation using Stanford HIVdb, while codon-frequency analysis provides additional information on the presence and relative abundance of minority resistance-associated variants that may not be represented in the consensus sequence. Together, these approaches provide a more detailed characterisation of HIVDR within individual samples, although read-level variant frequencies should not be interpreted as direct estimates of viral haplotype frequencies.

NanoHIV-DR currently targets the HIV-1 protease and reverse transcriptase regions and has not yet been evaluated for integrase or other genomic regions. Furthermore, the analytical reliability and clinical significance of low-frequency resistance variants detected using ONT sequencing require careful consideration, particularly given the potential influence of sequencing errors, amplification bias and coverage variability. Appropriate variant-frequency thresholds will therefore require further validation. Nevertheless, successful application to independently generated public ONT data provides preliminary evidence of workflow portability beyond the dataset used during development. The modular architecture also provides a foundation for future extensions to additional HIV-1 genomic regions and clinically relevant resistance-associated mutations.

## Acknowledgements

We thank Samantha Campbell for making her code publicly available through GitHub, which we adapted with minor modifications to generate clinical reports from Stanford HIVdb outputs.

